# The common *TMEM173 HAQ, AQ* alleles rescue CD4 T cellpenia, restore T-regs, and prevent *SAVI (N153S)* inflammatory disease in mice

**DOI:** 10.1101/2023.10.05.561109

**Authors:** Alexandra Aybar-Torres, Lennon A Saldarriaga, Ann T. Pham, Amir M. Emtiazjoo, Ashish K Sharma, Andrew J. Bryant, Lei Jin

## Abstract

The significance of STING (encoded by the *TMEM173* gene) in tissue inflammation and cancer immunotherapy has been increasingly recognized. Intriguingly, common human *TMEM173* alleles R71H-G230A-R293Q (*HAQ)* and G230A-R293Q (*AQ*) are carried by ∼60% of East Asians and ∼40% of Africans, respectively. Here, we examine the modulatory effects of *HAQ, AQ* alleles on STING-associated vasculopathy with onset in infancy (SAVI), an autosomal dominant, fatal inflammatory disease caused by gain-of-function human *STING* mutations. CD4 T cellpenia is evident in SAVI patients and mouse models. Using STING knock-in mice expressing common human *TMEM173* alleles *HAQ*, *AQ*, and *Q293*, we found that *HAQ, AQ*, and *Q293* splenocytes resist STING-mediated cell death *ex vivo,* establishing a critical role of STING residue 293 in cell death. The *HAQ/SAVI(N153S)* and *AQ/SAVI(N153S)* mice did not have CD4 T cellpenia. The *HAQ/SAVI(N153S), AQ/SAVI(N153S)* mice have more (∼10-fold, ∼20-fold, respectively) T-regs than *WT/SAVI(N153S)* mice. Remarkably, while they have comparable TBK1, IRF3, and NFκB activation as the *WT/SAVI*, the *AQ/SAVI* mice have no tissue inflammation, regular body weight, and normal lifespan. We propose that STING activation promotes tissue inflammation by depleting T-regs cells *in vivo*. Billions of modern humans have the dominant *HAQ, AQ* alleles. STING research and STING-targeting immunotherapy should consider *TMEM173* heterogeneity in humans.

**Teaser:** Common human *HAQ, AQ TMEM173* alleles dominate the gain-of-function *SAVI(N154S) TMEM173* mutant in mice.

## Introduction

STING drives cytosolic DNA-induced type I IFNs production ^1^. Recent research revealed that STING promotes inflammation in a variety of inflammatory diseases, including nonalcoholic fatty liver disease, nonalcoholic steatohepatitis, kidney injury, neurodegenerative diseases, cardiovascular diseases, obesity, diabetes, and aging ^2–9^. The type I IFNs-independent function of STING has also emerged ^9–12^. For example, initially described as a type I interferonopathy ^13^, recent studies in STING-associated vasculopathy with onset in infancy (SAVI) mouse models showed that SAVI is largely independent of type I IFNs ^14–17^. In a *STING N153S* mouse model of SAVI, crossing *N153S* mice to IRF3/IRF7, and IFNAR1 knockout mice, N153S mice still developed spontaneous lung diseases ^14^. JAK inhibitors were used to block type I IFNs signaling for SAVI patients with mixed success ^17–20^. For example, in a review of JAK inhibition in 18 SAVI patients, incomplete response to treatment happened in 7/18 (38%) of patients ^21^. Furthermore, 2 patients died of respiratory failure despite this treatment ^21^. Both radioresistant parenchymal and/or stromal cells and hematopoietic cells influence SAVI pathology in mice ^22,23^. The observation is important because it predicts that allogeneic stem cell transplantation may not work in human SAVI patients. Indeed, lung transplantations did not show improvement in SAVI patients ^18,24^. Patients died at 3- and 9 months post-lung transplant ^18,24^. How STING drives inflammation *in vivo,* independent of type I IFNs, remains unknown. Consequently, there is no curative care for SAVI.

Characterized as an innate immune sensor, STING expression is, paradoxically, high in CD4 T cells ^13,25^. Furthermore, STING activation kills mouse and human CD4 T cells *ex vivo* ^26–28^. SAVI patients and mouse models had CD4 T cellpenia ^13,28^. STING was first discovered as MPYS for its cell growth inhibition and cell death function in mouse B lymphoma cells ^29^. STING-mediated cell death is cell type dependent. For example, while STING activation kills human endothelial cells, primary and cancerous T cells, it does not kill mouse MEFs, BMDCs, or BMDMs ^30–32^. Second, STING-mediated cell death is type I IFNs-independent ^28,30,32^. Multiple cell death pathways, *i.e.,* apoptosis, necroptosis, pyroptosis, ferroptosis, and PANoptosis, are proposed ^32–34^. Last, the *in vivo* biological significance of STING-mediated CD4 T cell death is not clear ^28,32^. In humans, SAVI patients with constitutively activated STING have low CD4 T cell numbers ^13^, and type I IFNs are dispensable for STING-mediated human CD4 T cell death ^28^. Different from SAVI mice, SAVI patients (*N154S* or *V155M*) had normal counts of CD8 T and B cells ^13^.

The human *TMEM173* gene is highly heterogeneous ^35,36^. ∼50% of people in the U.S. carry at least one copy of non-*WT TMEM173* allele ^35^. Among them, the R71H-G230A-R293Q (*HAQ*) is the 2^nd^ most common *TMEM173* allele carried by ∼23% of people in the U.S. ^35^. However, in East Asians, *WT/HAQ* (34.3%), not *WT/WT* (22.0%), is the most common *TMEM173* genotype ^36^. Critically, the *HAQ* allele was positively selected in modern humans outside Africa ^37^. Anatomically modern humans outside Africa are descendants of a single Out-of-Africa Migration 50,000∼70,000 years ago. ∼1.4% of Africans have the *HAQ* allele, while ∼63.9% of East Asians are *HAQ* carriers ^37^. Haplotype analysis revealed that *HAQ* was derived from G230A-R293Q (*AQ*) allele ^37^. Importantly, the *AQ* allele was negatively selected outside Africa ^37^. ∼40.1% of Africans are *AQ* carriers, while ∼0.4% of East Asians have the *AQ* allele ^37^. *TMEM173* alleles often have a dominant negative effect likely because the protein STING exists as a homodimer ^29,38^. SAVI is an autosomal dominant inflammatory disease ^13^. *WT/HAQ* individuals had reduced Pneumovax23®-induced antibody responses compared to *WT/WT* individuals (NCT02471014) ^39^. Notably, *AQ* responds to CDNs and produces type I IFNs *in vivo* and *in vitro* ^37,40,41^, but the *AQ* allele was negatively selected in non-Africans ^37^. In contrast, the *HAQ* allele, defective in CDNs-type I IFNs responses ^35,36,39,42,43^, was positively selected in non-Africans ^37^, indicating that the CDNs-type I IFNs independent function of STING was essential for the survival of early modern humans outside of Africa.

In this study, we discovered, surprisingly, that the *HAQ*, *AQ* splenocytes are resistant to STING-mediated cell death. We generated *HAQ/SAVI(N153S)* and *AQ/SAVI(N153S)* mice and found that the *HAQ, AQ* alleles prevent CD4 T cellpenia, increasing/restoring T-regs and alleviating/stopping tissue inflammation in SAVI mice, thus providing evidence for the *in vivo* significance of type I IFNs-independent, STING-mediated cell death and potential *AQ*-based curative therapy for SAVI patients.

## Results

### STING activation kills mouse spleen CD4, CD8 T, and CD19 B cells *ex vivo*

We first used the synthetic non-CDNs STING agonist diABZI ^44^ to induce lymphocyte death because diABZI induces cell death without the need for lipid transfection or detergent for cell permeabilization ^31,45^ and diABZI is in clinical trials (NCT05514717). Splenocytes from C57BL/6N mice were treated with diABZI in culture, and cell death was determined by Annexin V and Propidium Iodide stain. Splenocyte cell death could be detected as early as 5 hours post diABZI treatment (Figure S1A). Dosage responses showed that ∼25ng/ml diABZI could kill 70% of splenocytes (Figure S1B). Similarly, STING agonists DMXAA and synthetic CDNs RpRpss-Cyclic di-AMP killed mouse spleen CD4, CD8 T cells, and CD19 B cells (Figure 1A). Thus, STING activation readily induces mouse lymphocyte death *ex vivo*.

**Fig. 1.**
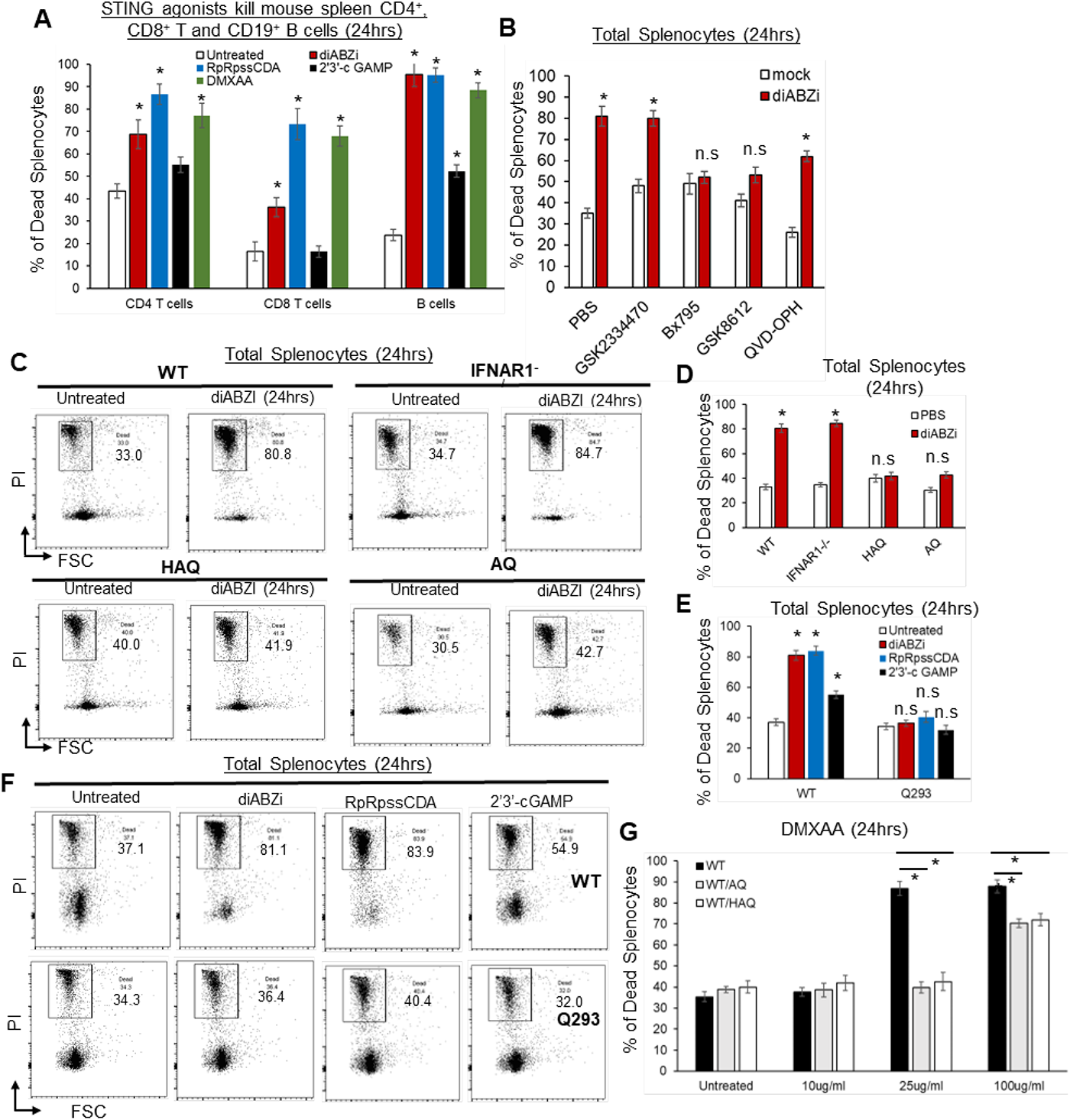
Splenocytes from *HAQ, AQ*, and *Q293* mice are resistant to STING-mediated cell death *ex vivo*. **(A)**. C57BL/6N splenocytes were treated directly (no transfection) with diABZI (100ng/ml), RpRpss-Cyclic di-AMP (5 μg/ml) or 2′3′-cGAMP (10μg/ml), DMXAA (25μg/ml) for 24hrs in culture. CD4, CD8 T cells and CD19 B cells death were determined by PI staining. (**B)**. Splenocytes from C57BL/6N mice were pre-treated with indicated small molecules, GSK2334470 (1.25µM), GSK8612 (2.5µM), Bx-795 (0.5µM), QVD-OPH (25µM) for 2hrs. diABZI (100ng/ml) was added in culture for another 24hrs. Dead cells were determined by PI staining. (**C-D).** Flowcytometry of *HAQ*, *AQ*, IFNAR1^-/-^ or C57BL/6N splenocytes treated with diABZI (100ng/ml) for 24hrs. Cell death was determined by PI staining. (**E-F)**. *Q293* or the *WT* littermates splenocytes were treated with diABZI (100ng/ml), RpRpss-Cyclic di-AMP (5 μg/ml) or 2′3′-cGAMP 10μg/ml) for 24h. Cell death was determined by PI staining. (**G).** *WT/HAQ*, *WT/AQ*, or *WT/WT* littermates splenocytes were treated with DMXAA (10, 25 or 100µg/ml) for 24h. Cell death was determined by PI staining. Data are representative of three independent experiments. Graphs represent the mean with error bars indication s.e.m. *p* values are determined by one-way ANOVA Tukey’s multiple comparison test (**A, E, G**) or unpaired student T-test (**B, D**). * p<0.05. n.s: not significant.

### TBK1 activation is required for STING-mediated mouse spleen cell death *ex vivo*

STING activation can lead to apoptosis, pyroptosis, necroptosis, or ferroptosis ^28–34,45,46^. We then treated mouse splenocytes with apoptosis, pyroptosis, necroptosis inhibitors, STING inhibitors H-151, C-176, and palmitoylation inhibitor 2-bromopalmitate (2-BP), followed by diABZI stimulation. Inhibitors for NLRP3 (MCC950), RIPK1 (Necrostatin-1), RIPK3 (GSK872), Caspase-1 (VX-795), Caspase-3 (Z-DEVD-FMK), Caspase 1,3,8,9 (Q-VD-Oph), ferroptosis (liproxstatin-1) did not affect diABZI-induced splenocyte cell death *ex vivo* (Figure S1B, S1C). The STING inhibitors H-151, C-176, and 2-BP also could not prevent diABZI-induced cell death (Figure S1C) though they inhibited diABZI-induced IFNβ production (Figure S1D). Instead, the TBK1 inhibitor BX-795 abolished diABZI-induced splenocyte death (Figure S1C).

BX-795 is a multi-kinase inhibitor, including 3-phosphoinositide-dependent protein kinase 1 (PDK1) and TBK1 (IC_50s_ = 6 and 11 nM, respectively). However, the treatment of PDK1 inhibitor GSK2334470 (IC50 = 10nM) did not prevent diABZI-induced splenocyte death (Figure 1B). In contrast, GSK8612, a highly potent and selective inhibitor for TBK1, prevented diABZI-induced splenocyte death (Figure 1B). Thus, TBK1 activation is likely critical for STING-mediated splenocyte cell death *ex vivo*.

### *HAQ, AQ, Q293 STING* knock-in mouse splenocytes are resistant to STING-mediated cell death *ex vivo*

*HAQ* and *AQ* are common human *TMEM173* alleles ^35–37^. Previously, we reported that *HAQ* knock-in mice are defective in CDNs-induced immune responses, while CDNs responses in *AQ* knock-in mice are similar to WT mice ^37^. We treated splenocytes from *HAQ* and *AQ* mice with diABZI *ex vivo* and found, surprisingly, that both *HAQ* and *AQ* splenocytes were resistant to diABZI-induced cell death (Figure 1C, 1D). In comparison, IFNAR1^-/-^ splenocytes were killed by diABZI, confirming that STING-mediated lymphocytes death are type I IFNs-independent (Figure 1C, 1D) ^28,30,32^.

*HAQ* and *AQ* share the common A230 and Q293 residues changes. We thus generated a *Q293 TMEM173* knock-in mouse. Notably, the Q293 splenocytes were resistant to STING agonists 2’3’-cGAMP, RpRpss-Cyclic di-AMP, and diABZI-induced cell death (Figure 1E, 1F). Thus, the residue 293 of STING is critical for its cell death function.

### *WT/HAQ*, *WT/AQ* mouse splenocytes are partially resistant to STING-mediated cell death *ex vivo*

*WT/HAQ* (34.3%) is the most common human *TMEM173* genotype in East Asians, while *WT/AQ* (28.2%) is the 2^nd^ most common *TMEM173* genotype in Africans ^36^. We generated *WT/HAQ*, *WT/AQ* mice and treated their splenocytes with mouse STING agonist DMXAA. *WT/HAQ* and *WT/AQ* splenocytes were protected from 25µg/ml DMXAA-induced cell death (Figure 1G). 100µg/ml DMXAA could kill *WT/HAQ* and *WT/AQ* splenocytes, albeit less than *WT/WT* cells (Figure 1G). Thus, the *HAQ* and *AQ* alleles are dominant and likely impact STING activation even in heterozygosity.

### STING activation kills primary human CD4 T cells but not CD8 T or CD19 B cells

STING agonists-based clinical trials in humans have been disappointing (NCT02675439, NCT03010176, NCT05514717) ^47,48^. We showed that the human *TMEM173* gene might undergo natural selection during the out-of-Africa migration ^37^ sensitive to evolutionary pressure. Thus, we investigated STING-mediated death in primary human lymphocytes.

Human explant lung cells from the *WT(R232)/WT(R232)* donors were treated with STING agonists 2’3-c GAMP, RpRpss-Cyclic di-AMP, diABZI for 24 hours in culture. Lymphocyte cell death was determined by Propidium Iodide staining. Different from mouse lymphocytes, diABZI and RpRpss-Cyclic di-AMP killed human CD4 T but not CD8 T or CD19 B cells (Figure 2A, 2B). Human CD8 T and CD19 B cells are resistant to 500ng/ml diABZI-induced cell death (Figure S2A).

**Fig. 2.**
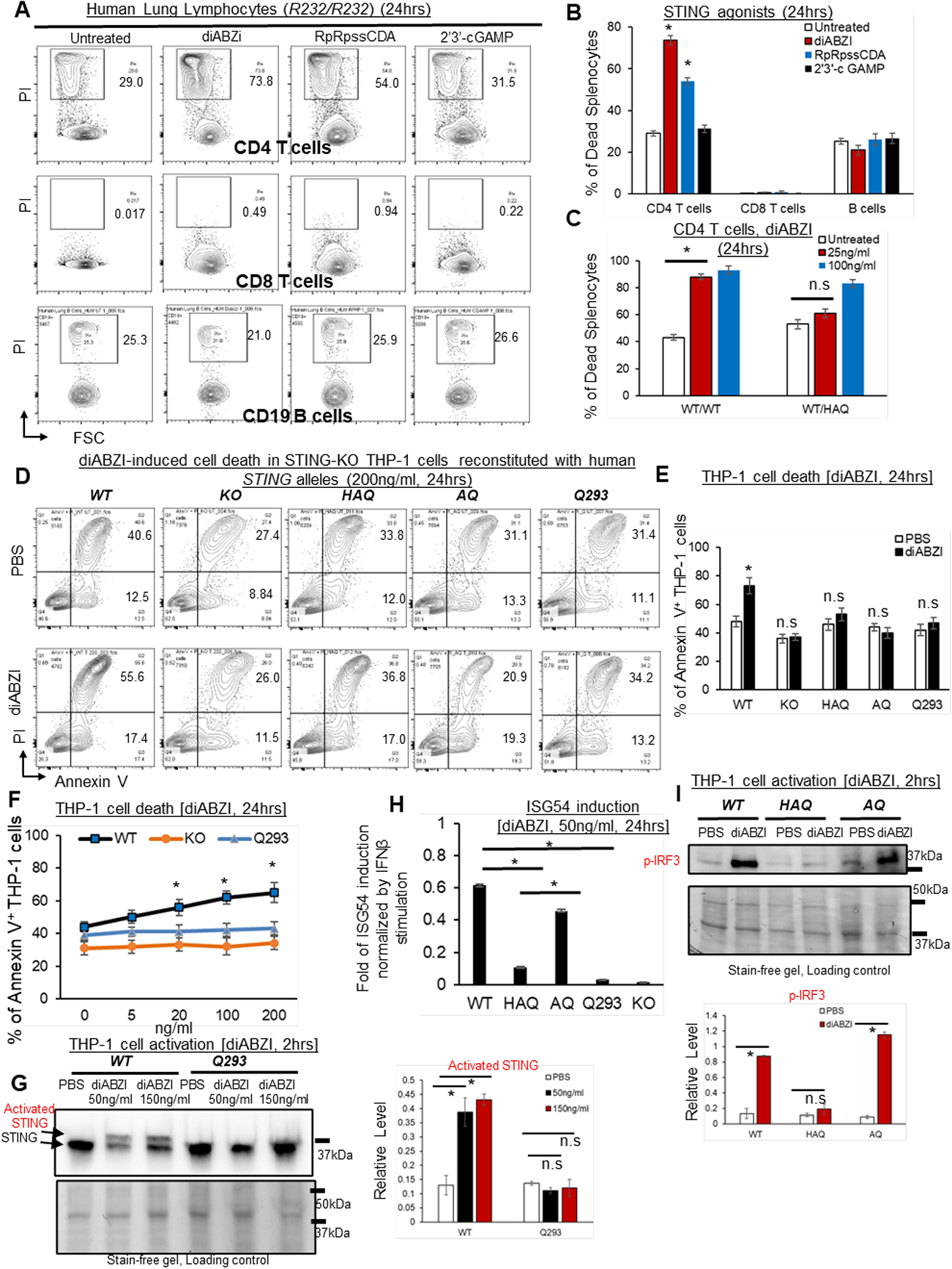
*HAQ*, *AQ*, and *Q293* human cells are resistant to STING agonists-induced death. **(A-B)**. Total human Lung cells from *WT/WT* individuals were treated with diABZi (100ng/ml) for 24hrs. Cell death in CD4, CD8 T cells and CD19 B cells were determined by PI staining. (**C)**. Total lung cells from a *WT/HAQ* (2 individuals) and a *WT/WT* (3 individuals) were treated with diABZi (25, 100ng/ml) for 24h. Cell death in CD4 T cells was determined by PI staining. (**D-E)**. STING-KO THP-1 cells (Invivogen,, cat no. thpd-kostg) were stably reconstituted with human *WT (R232), HAQ*, *AQ*, *Q293*. Cells were treated with diABZI (200ng/ml) in culture for 24 hrs. Dead cells were determined by Annexin V staining. (**F).** STING-KO THP-1 cells stably reconstituted with human *WT (R232), Q293* were treated with indicated dose of diABZI for 24hs in culture. Dead cells were determined by Annexin V staining. (**G**). STING-KO THP-1 cells stably reconstituted with human *WT (R232), Q293* were treated with indicated dose of diABZI for 2hs in culture. STING activation was detected by anti-STING antibody (Proteintech, #19851-1-AP). (**H)**. STING-KO THP-1 cells stably reconstituted with human *WT (R232), HAQ*, *AQ*, *Q293* were treated with 50ng/ml diABZI in culture for 24hrs. ISG-54 reporter luciferase activity was determined in cell supernatant and normalized to 10ng/ml IFNβ-stimulated ISG-54 luciferase activity. (**I).** STING-KO THP-1 cells stably reconstituted with human *WT (R232), HAQ*, *AQ* were treated with 50ng/ml diABZI in culture for 2hrs. STING and IRF3 activation were determined by anti-STING antibody and anti-p IRF3 antibody (CST, Ser396, clone 4D4G). Densitometry was determined by ImageLab 5. Data are representative of three independent experiments. Graphs represent the mean with error bars indication s.e.m. *p* values determined by one-way ANOVA Tukey’s multiple comparison test (**B, C, F, H, G**) or unpaired student T-test (**E, I**). * *p*<0.05, n.s: not significant.

### *WT/HAQ* human CD4 T cells are resistant to low doses of diABZI-induced cell death

*WT/HAQ* mouse splenocytes are resistant to low-dose diABZI-induced cell death (Figure 1G). To extend our observation into primary human T cells, we obtained lung explants from *WT/WT* and *WT/HAQ* individuals (Figure S2B) and treated them with diABZI in culture. 25ng/ml diABZI killed *WT/WT*, but not *WT/HAQ*, human lung CD4 T cells (Figure 2C).

### diABZI induces cell death in STING-KO human THP-1 cells reconstituted with WT human STING (R232) but not HAQ, AQ or Q293 human STING allele

To further determine cell death influenced by human *TMEM173* alleles *HAQ, AQ,* and *Q293*, we used the *STING-KO* THP-1 cell line because STING agonist induces type I IFNs and cell death in *STING-KO* THP-1 cells expressing *WT* human STING ^34^ (Figure S2C, S2D). We, thus, generated stable THP-1 *STING-KO* lines expressing *HAQ, AQ, WT,* or *Q293 TMEM173* allele. Cell death was determined by Annexin-V staining. diABZI killed THP-1 *STING-KO* lines expressing *WT* but not *HAQ, AQ,* or *Q293 TMEM173* allele (Figure 2D, 2E). No cell death was induced in the *Q293* THP-1 cells stimulated by 20 to 200 ng/ml of diABZI (Figure 2F). diABZI also did not induce STING activation in *Q293* THP-1 cells (Figure 2G). Notably, 50ng/ml diABZI induced p-IRF3 activation and type I IFNs in *AQ* THP-1 cells but not *HAQ* THP-1 cells (Figure 2H, 2I), indicating that the STING-cell death and STING-IRF3-Type I IFNs pathways can be uncoupled.

### *HAQ* and *AQ* alleles rescue the lymphopenia and suppress myeloid cell expansion in ***SAVI(N153S)* mice.**

The *in vivo* significance of the STING/MPYS-cell death is unclear. Furthermore, multiple cell death pathways, i.e., apoptosis, necroptosis, pyroptosis, ferroptosis, and PANoptosis, are proposed ^32–34^. The uncertainty likely results from studies using different cell types (primary cells vs cancer cell lines); species (human vs mouse); STING agonists (cGAMP, which requires cell permeabilization by detergents or lipid transfection, vs diABZi, DMXAA that can directly cross the membrane) ^26,27,30,31,49^. Critically, which mechanism is relevant *in vivo*, causing T cellpenia is not known. To clarify the *in vivo* significance and mechanisms of STING-mediated cell death, we turned to SAVI mice.

SAVI is an autosomal dominant, inflammatory disease caused by one copy of a gain-of-function *TMEM173* mutant (*WT/SAVI*) ^13^. CD4 T cellpenia was found in SAVI patients and SAVI mouse models ^13,14^. STING activation in SAVI mice is independent of ligands and happens *in vivo*. We thus generated *HAQ/SAVI(N153S)* and *AQ/SAVI(N153S)* mice aiming to establish the *in vivo* significance and mechanism of STING-cell death.

First, *HAQ/SAVI(N153S)* and *AQ/SAVI(N153S)* mice had reduced splenomegaly compared to *WT/SAVI(N153S)* mice though their spleens were still larger than the littermates *WT/HAQ* and *WT/AQ* (Figure 3A, 3B). Next, *HAQ/SAVI(N153S)* and *AQ/SAVI(N153S)* mice had similar spleen B cells and CD4 T cell numbers as the *WT/HAQ*, *WT/AQ* littermates (Figure 2B, 2C). Their CD8^+^ T cells were lower than their *WT* littermates but much higher than the *WT/SAVI(N153S)* mice (Figure 3D). Third, spleen myeloid cell numbers, i.e., neutrophils, Ly6C^hi^ monocytes and F4/80 macrophages, were all reduced by half compared to *WT/SAVI(N153S)* mice (Figure 3E∼3H). Last, splenocytes from *HAQ/SAVI*, *AQ/SAVI* mice were partially resistant to diABZI, DMXAA-induced cell death *ex vivo* (Figure S3). Notably, the *HAQ/SAVI(N153S)* and *AQ/SAVI(N153S)* mice also had restored bone marrow monocytes (Figure S4). Thus, *HAQ* and *AQ* alleles prevent lymphopenia and suppress myeloid cell expansion in *SAVI(N153S)* mice.

**Fig. 3.**
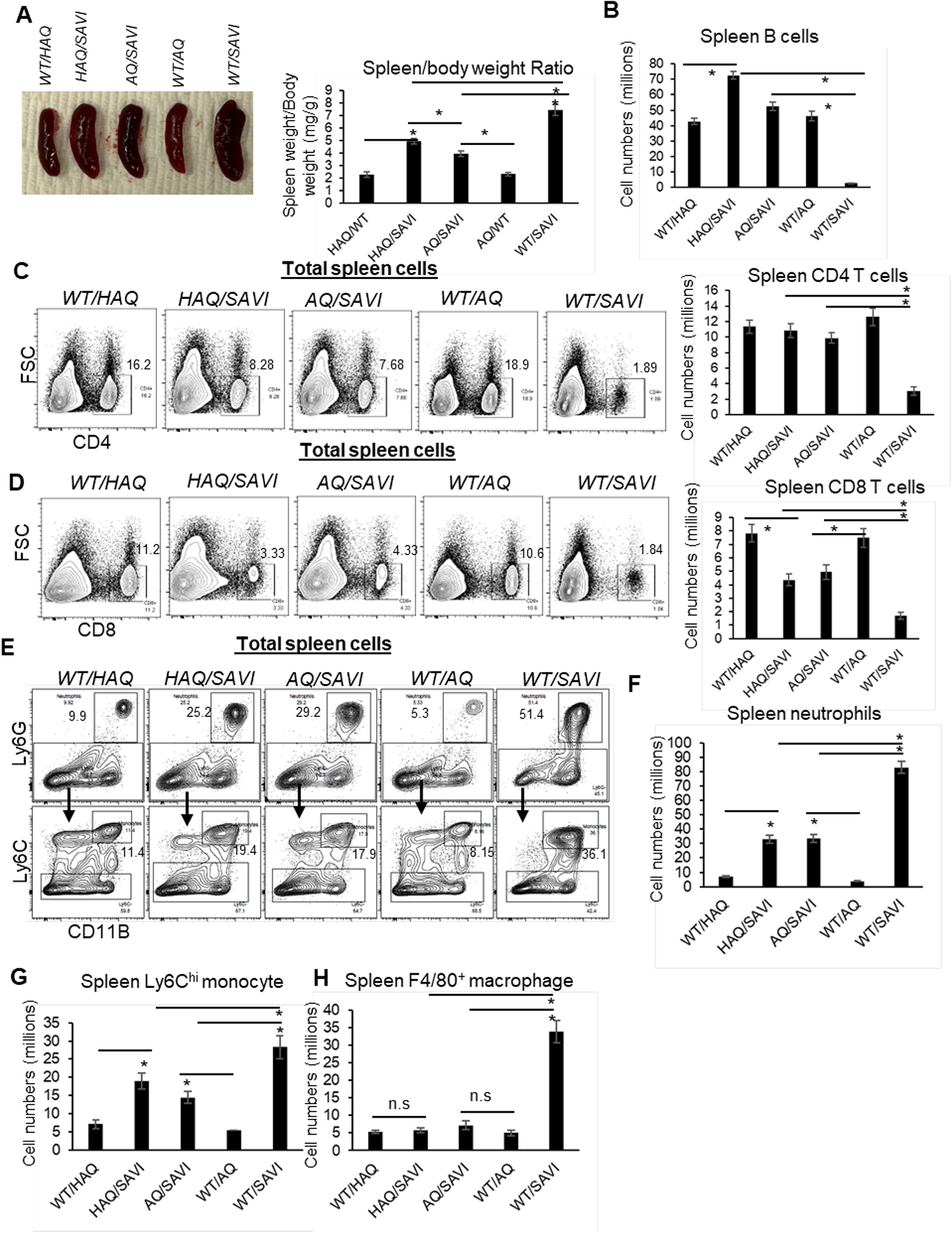
*HAQ* and *AQ* rescue the lymphopenia and suppress myeloid cell expansion in *SAVI (N153S)* mice. **(A)**. The size and weight of spleens from *WT/HAQ, HAQ/SAVI, AQ/SAVI, WT/AQ, WT/SAVI*. **(B-D).** Spleen CD19^+^ B cells, CD4, CD8 T cells were determined in the indicated mice by Flow. (**E-H).** Spleen Ly6G^+^ neutrophils, Ly6C^hi^ monocytes and F4/80^+^ macrophage was determined in the indicated mice by Flow. Data are representative of three independent experiments. n=3∼5 mice/group. Graphs represent the mean with error bars indication s.e.m. *P* values are determined by one-way ANOVA Tukey’s multiple comparison test. * *p*<0.05, n.s: not significant.

### The *HAQ* allele alleviates and the *AQ* allele prevents *SAVI(N153S)* disease in mice

*SAVI(N153S)* disease is characterized by early onset, failure to thrive (low body weight), persistent lung inflammation, decreased lung function, and young death in humans and mouse models ^13,17,49,50^. The *HAQ/SAVI(N153S)* mice weighed more and had an improved lifespan than the *WT/SAVI(N153S)* mice (Figure 4A, 4B). The lifespan, airway resistance, and tissue inflammation (lung, liver) were also improved in *HAQ/SAVI(N153S)* mice compared to the *WT/SAVI(N153S)* mice (Figure 4C, 4D, 4J, 4K). However, the pulmonary artery pressure was still elevated in *HAQ/SAVI(N153S)* mice (Figure 4E). Remarkably, the *AQ/SAVI(N153S)* mice had similar body weight and lifespan as the *WT/AQ* mice (Figure 4F, 4G). The airway resistance, pulmonary artery pressure, and tissue inflammation in *AQ/SAVI(N153S)* were similar to the *WT/AQ* littermates (Figure 4H, 4I, 4J, 4K). Thus, the *HAQ* allele alleviates and the *AQ* allele prevents inflammatory SAVI disease in mice. Interestingly, lungs from *HAQ/SAVI, AQ/SAVI* had similarly reduced infiltration of Ly6G^+^CD11B^+^ neutrophils and Ly6G^-^Ly6C^+^CD11B^+^ inflammatory monocytes compared to *WT/SAVI* mice (Figure S5).

**Fig. 4.**
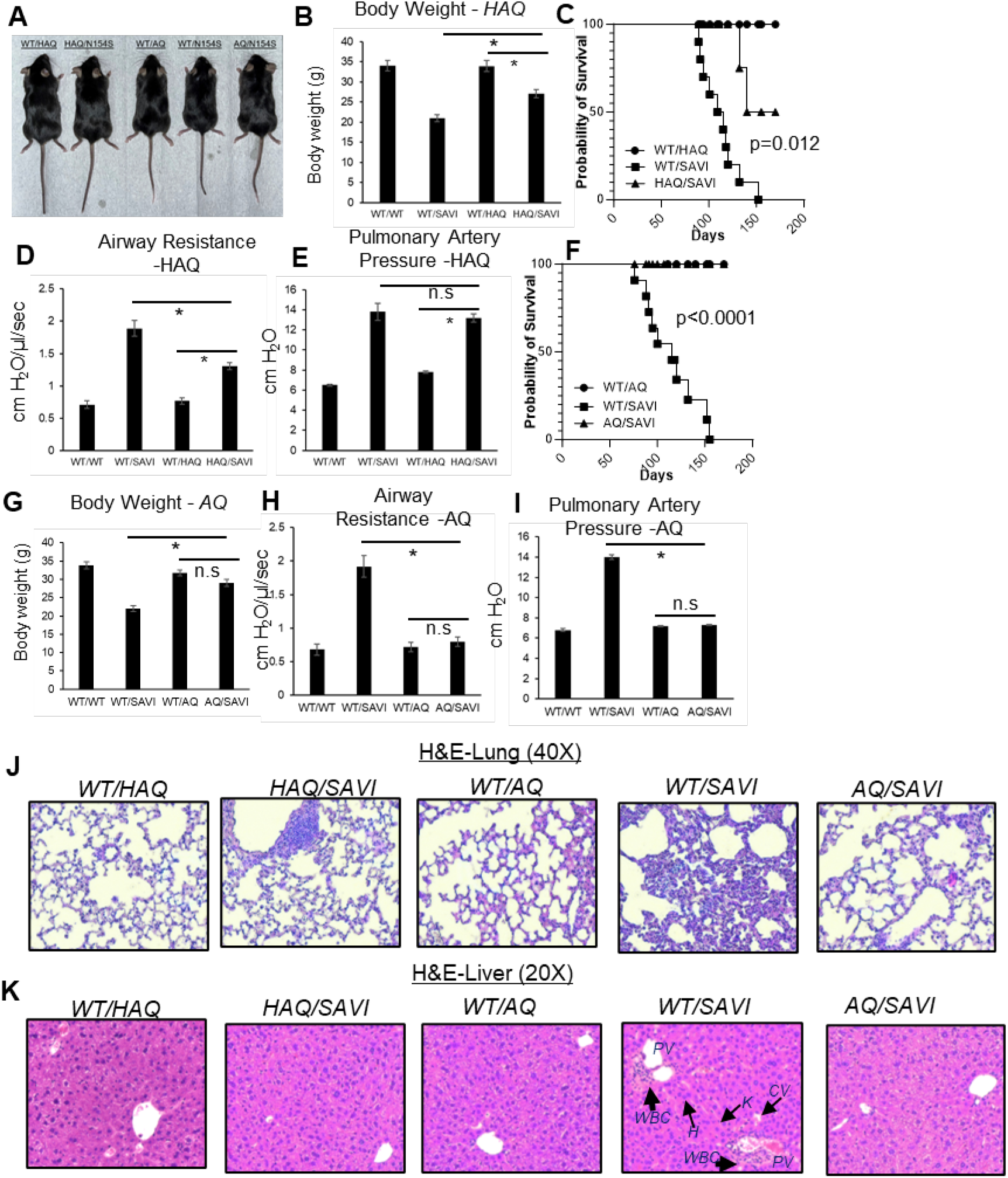
*HAQ* and *AQ* alleles prevent *SAVI(N153S)* disease in mice. **(A, B, G)**. The size and body weight of *HAQ/SAVI*, *AQ/SAVI*, *WT/SAVI* and their littermates *WT/HAQ*, *WT/AQ* mice. (**D, E, H, I)**. Airway resistance, and pulmonary artery pressure were determined as described in **Materials and Methods**. **(C, F).** *HAQ/SAVI, AQ/SAVI, WT/SAVI* (10 mice/group), were monitored for survival by Kaplan-Meier. (**J, K**). Representative hematoxylin and eosin (H&E) staining of lung, liver sections from indicated mice. n=3∼5 mice/group. Data are representative of 3 independent experiments. Graphs represent the mean with error bars indication s.e.m. *p* values are determined by one-way ANOVA Tukey’s multiple comparison test. * *p*<0.05, \*\**p*<0.01. n.s.: not significant. **WBC**: white blood cells; **H**: hepatocytes; **K**: Kupper cells; **PV**: portal vein; **CV**: central vein.

### diABZI induces similar STING, TBK1, IRF3, NFκB activation in the *AQ/SAVI(N153S)* and *WT/SAVI(N153S)* bone-marrow-derived-macrophage (BMDM)

SAVI was characterized as type I interferonopathy ^17^. However, several studies showed that type I IFN signaling and IRF3 activation were dispensable for SAVI disease ^14,15,23,50^. *AQ* allele prevents SAVI disease (Figure 4). However, diABZI-treated *AQ/SAVI(N153S)* and *WT/SAVI(N153S)* BMDM had similar TBK1-IRF3 activation and IFNβ production (Figure 5A, 5B). diABZI treatment caused IκBα degradation, and similar TNF production in *WT/SAVI* and *AQ/SAVI* BMDM (Figure 5C, 5D). Furthermore, diABZI activation led to STING protein degradation in *WT/SAVI* and *AQ/SAVI* BMDM (Figure 5E). Last, using cleavable crosslinker dithiobis (succinimidyl propionate (DSP), we showed that STING in *WT/SAVI, AQ/SAVI* BMDM forms a similar dimer *in situ* (Figure 5F). Thus, the *AQ/SAVI* BMDM had similar STING degradation, TBK1, IRF3, NFκB activation, and dimerization as the *WT/SAVI* BMDM.

**Fig. 5.**
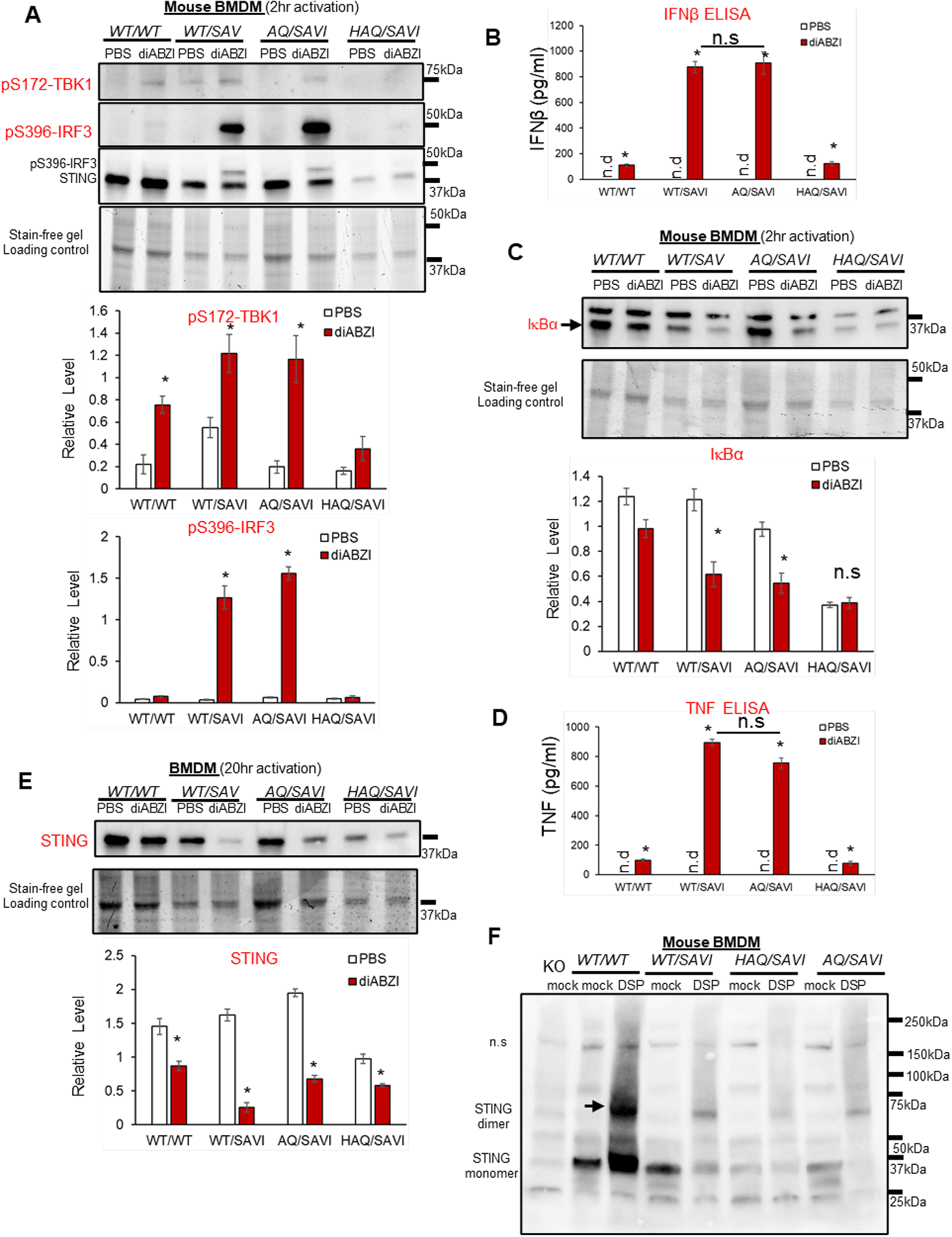
*AQ/SAVI(N153S)* cells had similar TBK1-IRF3, NFκB activation and STING degradation as the *WT/SAVI(N153S)* cells. (**A, C).** BMDM from *WT/WT, WT/SAVI, HAQ/SAVI* and *AQ/SAVI* were treated with 100ng/ml diABZi in culture for 2 hrs. Cells were lysed and run on a 4∼20% Mini-PROTEAN TGX Stain-Free Precast Gel. The blot was probed for phospho-Thr172-TBK1 antibody (CST, clone D52C2), phosphor-Ser 396-IRF3 (CST, clone 4D4G), STING (Proteintech, #19851-1-AP) and IκBα (CST, clone 44D4) antibody. **(E).** BMDM from *WT/WT, WT/SAVI, HAQ/SAVI* and *AQ/SAVI* were treated with 100ng/ml diABZi in culture for 24 hrs. Cells were lysed and run on a 4∼20% Mini-PROTEAN TGX Stain-Free Precast Gel. The blot was probed for STING antibody (Proteintech, 19851-1-AP). (**B, D**). IFNβ and TNF were determined by ELISA in the cell supernatant from **E**. (**F).** BMDM from *WT/WT, WT/SAVI, HAQ/SAVI* and *AQ/SAVI* were treated with 400µM cleavable chemical crosslinker DSP (Pierce™, cat no: PG82081) in PBS for 1 hr at 4°C. Cells were washed with PBS and lysed in RIPA buffer. Whole cell lysate was mixed with 4x Laemmli Sample Buffer (BioRad, cat no 1610747) containing 5% 2-mercaptoethanol, heated at 95°C for 10 min and, run on a 4∼20% Mini-PROTEAN TGX Stain-Free Precast Gel. The blot was probed for STING antibody (Proteintech, 19851-1-AP). Densitometry was determined by ImageLab 5. Data are representative of three independent experiments. Graphs represent the mean with error bars indication s.e.m. *p* values are determined by unpaired student T-test (**A-E**) or one-way ANOVA Tukey’s multiple comparison test (**D, B**). * *p*<0.05, \*\**p*<0.01, \*\*\**p*<0.001. n.s.: not significant; n.d: not detected.

### *The HAQ* allele increased, and the *AQ* allele restored T-regs in *SAVI(N153S)* mice

IFNγ was proposed to drive SAVI disease ^15,23,38^. We confirmed that *WT/SAVI* CD4 T cells were enriched with IFNγ^+^ cells (Figure 6A). However, *WT/SAVI* mice have CD4 T cellpenia. Thus, the total numbers of spleen IFNγ^+^ CD4 T cells were comparable in *WT/SAVI* and *AQ/SAVI* mice (Figure 6A). In contrast, the *HAQ/SAVI mice* had decreased IFNγ^+^ CD4 T cells (Figure 6A).

**Fig. 6.**
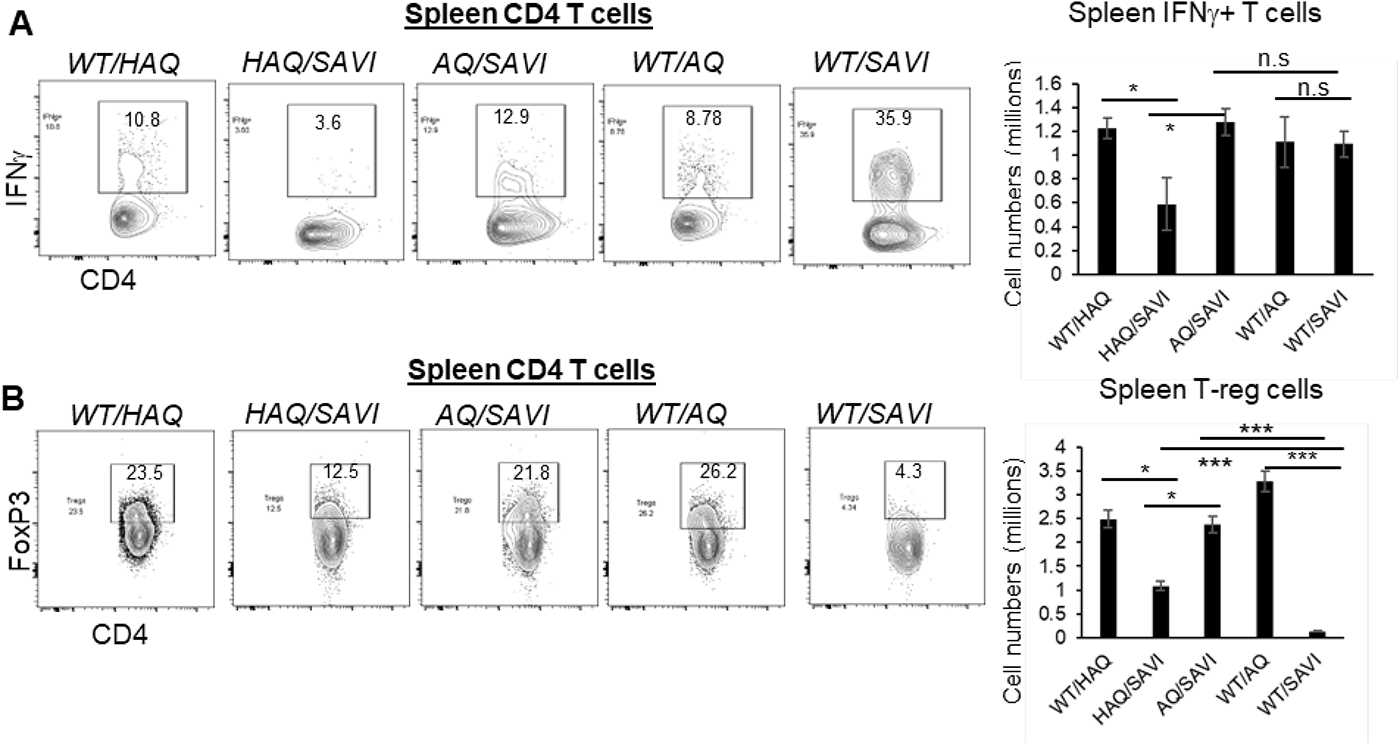
*HAQ/SAVI(N153S)* and *AQ/SAVI(N153S)* cells had 10-fold and 20-fold increased spleen T-regs compared to *WT/SAVI* mice. **(A)**. Flow cytometry analysis of IFNγ producing spleen CD4^+^ T cells from *WT/HAQ, WT/AQ, WT/SAVI, HAQ/SAVI* and *AQ/SAVI* mice. (**B)**. Flow cytometry analysis of CD4^+^ FoxP3^+^ spleen T-regs. n=3∼5 mice/group. Data are representative of 3 independent experiments. Graphs represent the mean with error bars indication s.e.m. *p* values are determined by one-way ANOVA Tukey’s multiple comparison test. * *p*<0.05, \*\**p*<0.01, \*\*\**p*<0.001. n.s.: not significant.

The induction of Foxp3 expression in T-reg cells during ongoing autoimmune inflammation resolved inflammation and pathology in mice ^51^. CD4 T cellpenia depletes CD4 T-regs. Indeed, *WT/SAVI* mice had ∼20-fold reduction of spleen FoxP3^+^ T-regs compared to *AQ/SAVI* or *WT/WT* littermates (Figure 6B). The *HAQ/SAVI* mice also had ∼10-fold more T-regs than the *WT/SAVI* littermate (Figure 6B).

## Discussion

This study, using the *HAQ, AQ, SAVI(N153S) TMEM173* knock-in mice, reveals the *in vivo* significance and mechanism of STING-mediated CD4 T cell death. *HAQ, AQ* alleles prevent CD4 T cellpenia, and increase/restore CD4 T-regs in SAVI mice. The results are consistent with previous finding that the impaired CD4 T cell proliferation by the *SAVI(V155M)* mutant could be rescued by the addition of the *HAQ* allele *in vitro* ^27^. STING has been increasingly implicated in inflammatory diseases such as nonalcoholic fatty liver disease, nonalcoholic steatohepatitis, cardiomyopathy, obesity, diabetes, neurodegenerative diseases, aging, and kidney injury, many of which are independent of type I IFNs ^2,3,9^. It is tempting to suggest that STING activation in CD4 T cells leads to CD4 T-regs depletion that break tissue tolerance and exacerbates tissue inflammation.

Human immunodeficiency virus (HIV) primarily infects CD4 T cells and might activate the STING pathway in CD4 T cells ^52–57^. The loss of CD4 T cells is the hallmark of untreated HIV infection ^58,59^, and the measurement of CD4 T cell count is a central part of HIV care. We found that *HAQ* and *AQ* CD4 T cells are resistant to STING-mediated cell death. Mogensen and colleagues reported that *HAQ/HAQ* was enriched in HIV-infected long-term nonprogressors ^42^. These *HAQ/HAQ* individuals had reduced inhibition of CD4 T cell proliferation and a reduced immune response to DNA and HIV ^42^. It is likely that HIV infection activates STING-cell death pathway in CD4 T cells. In *HAQ/SAVI* and *AQ/SAVI* mice, one copy of *HAQ*, *AQ* allele suppressed CD4 T cell death. *HAQ*, *AQ* carriers might have fewer HIV-induced CD4 T cell death, thus being long-term nonprogressors in HIV infection-induced Acquired immunodeficiency syndrome (AIDS)^42^. Targeting STING to prevent CD4 T cell death might be a valid therapy for AIDS. Activating the STING pathway is a promising strategy for cancer immunotherapy ^60–66^.

Multiple STING agonists are in clinical trials ^47,48^. Recently, the safety issue emerged in some STING agonist trials ^47,48^. For example, in the STING agonist, ADU-S100, clinical trial, Grade 3/4 treatment-related adverse events were reported in 12.2% of 41 pretreated patients (NCT02675439)^47,48^. The National Institutes of Health defines grade 3 as “incapacitating; unable to perform usual activities; requires absenteeism or bed rest.” In a clinical trial using STING antibody-drug-conjugate (ADC) that conjugates diABZI to anti-HER2 Ab, a Grade 5 (fatal) serious adverse event was recorded and deemed related to the STING-ADC (NCT05514717). SAVI disease, driven by overreacting STING, is often fatal^13^. *AQ*, to a less degree, *HAQ*, suppress mortality in SAVI mice. Future STING clinical trials should be based on human *TMEM173* genotype to achieve safe and effective responses.

Mechanistically, apoptosis, pyroptosis, ferroptosis, necroptosis, and PANoptosis have all been reported in STING-mediated cell death ^28–34,45,46^. Different cell types and STING agonists used likely contributed to the inconsistency and complexity. Here, we focused on lymphopenia in the SAVI mice that avoids ligand-dependent, non-physiological dosage in STING-mediated cell death. *HAQ* and *AQ* alleles could prevent CD4 T cellpenia in the SAVI mice strongly indicating that residue A230 or Q293 prevent STING-mediated CD4 T cell death *in vivo*. Splenocyte from *Q293* mice were resistant to STING agonists-induced cell death *ex vivo*. Thus, it is likely that the Q293 residue is critical for STING-mediate lymphopenia. Notably, Q293 is outside the C-terminal tail (CTT) (residues 341–379 of human STING) critical for TBK1 recruitment and IRF3 phosphorylation ^67^ or miniCTT domain (aa343–354) ^27^, or the UPR motif (aa322–343) ^49^ important for T cells death *in vitro*. Further studies are needed to understand how the aa293 of STING mediates cell death *in vivo*. Noteworthy, *AQ/SAVI* cells had similar TBK1-IRF3, NFκB activation and STING degradation as the *WT/SAVI* cells. Yet, *AQ/SAVI* mice did not have CD4 T cellpenia as *WT/SAVI* mice suggesting that the canonical STING-TBK1-IRF3/NFκB pathway, likely STING oligomerization, is not sufficient for the induction of cell death at the physiological condition.

We used the *WT/N153S* knock-in SAVI mouse model that spontaneously develop lung inflammation, T cell cytopenia, and early mortality, mimicking pathological findings in human SAVI patients^16^. Using the *WT/N153S* SAVI mouse model and human Jurkat T cell line, it was proposed that STING activation causes chronic ER stress and unfolded protein response, leading to T cell death by apoptosis ^49^. Furthermore, the study showed that crossing *WT/N153S* mice to the OT-I mice reduced ER stress and restored CD8^+^, but not CD4^+^, T cells ^49^. The restoration of CD8^+^T cells reduces inflammation and lung disease ^49^. However, human *WT/N154S* SAVI patients have normal CD8^+^ T cells numbers ^13^, and primary human CD8^+^ T cells are largely resistant to STING-agonists-induced cell death *ex vivo* (Figure 2A) ^28^. Thus, it is puzzling how restoring CD8^+^ T cells can rescue SAVI phenotypes since the SAVI patients already have normal CD8^+^ T cells numbers.

Finally, it is unexpected that both *HAQ* and *AQ* alleles are resistant to cell death. Our previous studies showed that the *HAQ* and *AQ* alleles have opposite functions ^37^. AQ-STING, not HAQ-STING, responds to CDNs ^35–37,39–43,68^. *AQ* mice are lean while *HAQ* mice are fat ^37^. Most importantly, *HAQ* was positively selected, while *AQ* was negatively selected, in modern humans outside Africans ^37^. Thus, the death pathway of STING is also distinct from the STING function that was naturally selected. It is worth noting that the *AQ* allele does better than the *HAQ* allele in suppressing SAVI disease. Thus, besides preventing cell death, additional mechanism by the *AQ* allele, e.g. fatty acid metabolism ^37,69^, is involved in curing SAVI.

### The limitations of the study

The poor transferability of mouse to humans is a major issue in STING research ^47,48^. The present study used *AQ/SAVI* and *HAQ/SAVI* mice. Confirmation is needed in humans with the identification and evaluation of people who are *AQ/SAVI*, *HAQ/SAVI*.

## Materials and Methods

### Experimental Design

The study was designed to reveal (i) the *in vivo* significance of the type I IFNs-independent, STING-dependent cell death function; (ii) the interplay between common *STING* alleles *HAQ, AQ* and the rare, gain-of-function SAVI STING mutation; (iii) the driver for the inflammatory SAVI disease. Mouse splenocytes, primary human lung cells, human THP-1 cells and *HAQ, AQ, SAVI* knock-in mice were used to establish the *in vivo* significance and human relevance. All the repeats were biological replications that involve the same experimental procedures on different mice. Where possible, treatments were assigned blindly to the experimenter by another individual in the lab. When comparing samples from different groups, samples from each group were analyzed in concert, thereby preventing any biases that might arise from analyzing individual treatments on different days. All experiments were repeated at least twice.

### Mice

*WT/SAVI(N153S)* mice were purchased from The Jackson Laboratory. *HAQ*, *AQ* mice were previously generated in the lab ^36,37^. The *Q293* mice were generated by Cyagen Biosciences. Briefly, the linearized targeting vector was transfected into JM8A3.N1 C57BL/6N embryonic stem cells. A positive embryonic stem clone was subjected to the generation of chimera mice by injection using C57BL/6J blastocysts as the host. Successful germline transmission was confirmed by PCR sequencing. The heterozygous mice were bred to Actin-flpase mice [The Jackson Laboratory, B6.Cg-Tg (ACTFLPe)9205Dym/J] to remove the neo gene and make the *Q293* knock-in mouse. Age-and gender-matched mice (2 – 6 month old, both male and female) were used for indicated experiments. *WT/SAVI* (male) x *WT/HAQ* (female), *WT/SAVI* (male) x *WT/AQ* (female) breeders were set up to generate *HAQ/SAVI*, *AQ/SAVI* mice. Mice were housed at 22°C under a 12-h light-dark cycle with ad libitum access to water and a chow diet (3.1 kcal/g, Teklad 2018, Envigo, Sommerset, NJ) and bred under pathogen-free conditions in the Animal Research Facility at the University of Florida. Littermates of the same sex were randomly assigned to experimental groups. All mouse experiments were performed by the regulations and approval of the Institutional Animal Care and Use Committee at the University of Florida, IACUC202200000058.

### Reagents

Recombinant human IFNβ (R&D, cat no. 8499-IF-010/CF), diABZI (Invivogen, cat no. 2138299-34-8), 2’3’-cGAMP (Invivogen, cat no. tlrl-nacga23-02), DMXAA (Invivogen, cat no. tlrl-dmx), H151 (Invivogen, cat no. inh-h151), RpRpSS-Cyclic di-AMP (Biolog, cat no. C118), THP1-Dual™ KO-STING Cells (Invivogen, cat no. thpd-kostg). All other chemical inhibitors are from Selleckchem. Mouse TNF alpha ELISA Ready Set Go. (eBioscience, cat no. 88-7324). Mouse IFN-Beta ELISA Kit (PBI, cat no. 42400).

### Generation of THP-1 KO-STING cells stably expressing human *TMEM173* alleles

THP1-Dual™ KO-STING Cells in 6-well plate were transfected with 1µg *TMEM173* plasmid (in pcDNA 3.1 vector) with Lipofectamine LTX and Plus Reagent (Invitrogen, cat no: A12621) according to the manufacturer’s instructions. <u>Transfecting Plasmid DNA into THP-1 Cells Using Lipofectamine LTX Reagent | Thermo Fisher Scientific - US</u>. 48hrs after the transfection, the cell medium was changed. G418 (1mg/ml) was added to the culture to select STING expressing THP-1 cells. The G418-resistant cells were established and expanded.

### Histology

Lungs and livers were fixed in 10% formalin, paraffin-embedded, and cut into 4-µm sections. Lung, liver sections were then stained for hematoxylin-eosin. All staining procedures were performed by the histology core at the University of Florida. Briefly, tissue sectins were immersed Harris Hematoxylin for 10 seconds, then washed with tap water. Cleard sections were re-immersed in EOSIN stain for ∼30 seconds. The sections were washed with tap water until clear, then dehydrate in ascending alcohol solutions (50%,70%,80%,95% x 2, 100% x 2). Afterwards, the sections werer cleared with xylene (3 - 4 x). The sections were mounted on glass slide with permount organic mounting medium for visulization.

### Lung Function

Pulmonary function was evaluated using an isolated, buffer-perfused mouse lung apparatus (Hugo Sachs Elektronik, March-Huggstetten, Germany), as previously described ^70^. Briefly, mice were anesthetized with ketamine and xylazine and a tracheostomy was performed, and animals were ventilated with room air at 100 breaths/min at a tidal volume of 7 μl/g body weight with a positive end-expiratory pressure of 2 cm H2O using a pressure-controlled ventilator (Hugo Sachs Elektronik, March-Huggstetten, Germany).

### Isolation of lung cells

Cells were isolated from the lung as previously described ^71^. The lungs were perfused with ice-cold PBS and removed. Lungs were digested in DMEM containing 200μg/ml DNase I (Roche, 10104159001), 25μg/ml Liberase TM (Roche, 05401119001) at 37°C for 2 hours. Red blood cells were then lysed and a single cell suspension was prepared by filtering through a 70-µm cell strainer.

### BMDM activation

BMDMs were induced from mouse bone marrow cells cultured in RPMI 1640 (cat#11965; Invitrogen) with 10% FBS, 2 mM L-glutamine, 1 mM sodium pyruvate, 10 mM HEPES buffer, 1% nonessential amino acids, 50 mM 2-ME, 1% Pen/Strep, with 20ng/ml M-CSF (Kingfisher, RP0407M) for 10 days ^36^. STING agonists were added into the culture (no transfection or membrane permeabilization).

### Flow cytometry

Single cell suspensions were stained with fluorescent-dye-conjugated antibodies in PBS containing 2% FBS and 1mM EDTA. Surface stains were performed at 4°C for 20 min. For intracellular cytokine or transcription factor staining of murine and human cells, cells were fixed and permeabilized with the Foxp3 staining buffer set (eBioscience, cat no 00-5523-00). Cells were washed and stained with surface markers. Cells were then fixed and permeabilized (eBioscience, cat no. 00-5523-00) for intracellular cytokine stain. Data were acquired on a BD LSRFortessa and analyzed using the FlowJo software package (FlowJo, LLC). Cell sorting was performed on the BD FACSAriaIII Flow Cytometer and Cell Sorter. More information on the antibodies used can be found in supplementary table S1.

### Human lung explants

Human lung explants were procured at the Lung Transplant Center, Division of Pulmonary, Critical Care and Sleep Medicine, Department of Medicine, University of Florida. Donor and patients consent was obtained for a research protocol (UF IRB201902955-Treatment with IFNβ Induces Tolerogenic Lung Dendritic Cells in Human advanced lung disease). Healthy donor lungs were surgically removed postmortem, perfused, small pieces were cut from the right middle and lower lobes for research purpose, and stored in cold Perfadex ® at 4°C for no more than 12 hrs before processing. Ex planted lungs from emphysema lung transplant patients were stored in cold Perfadex ® at 4°C for no more than 12 hrs before the process. No lung explants were procured from prisoners.

### Statistical Analysis

To gain statistical power, we employ three∼four mice/groups to characterize lung immunity. Ten mice/group to monitor animal health. The statistical justification for group size was calculated using the SAS program to calculate the animal numbers. The analysis was carried out using a standard error of 0.5 for immunological assays, and a power of 0.9. All data are expressed as means ± SEM. Statistical significance was evaluated using Prism 9.0 software. Comparisons between two groups were analyzed by performing an unpaired Student’s *t* test. Comparisons between more than two groups were analyzed by performing a one-way analysis of variance (ANOVA) with Tukey’s multiple comparisons test.

## Author contributions

Conceptualization: LJ

Methodology: AAT, LAS, ATP, AME

Investigation: AAT, LAS, ATP, AKS, AJB, LJ

Supervision: LJ

Writing—original draft: LJ

Writing—review & editing: LJ, AJB, AKS

## Data and materials availability

All data are available in the main text or the supplementary materials.

## Supplementary Materials

**Fig. S1.**
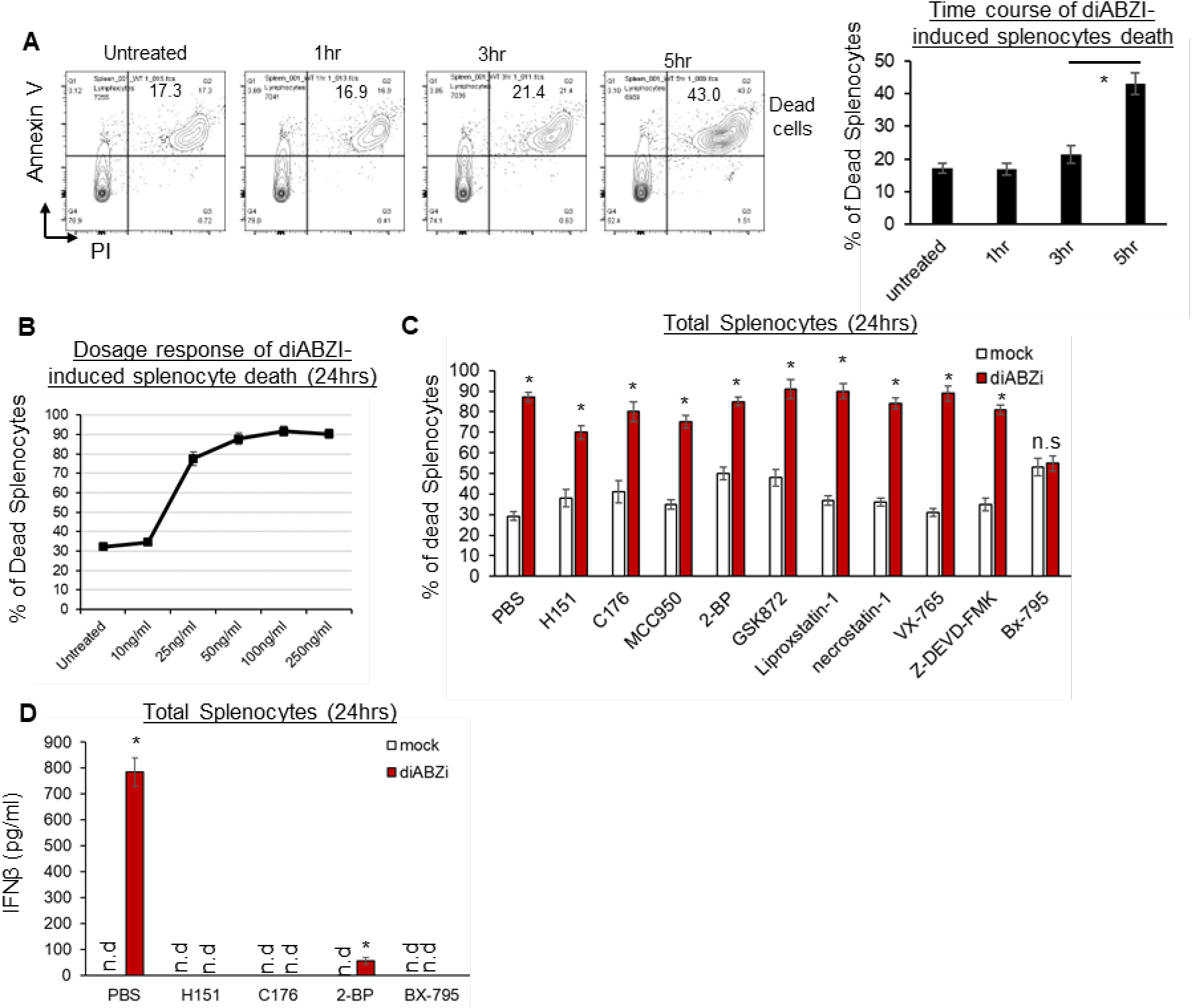
Bx-795 inhibits diABZI-induced mouse splenocyte death. **(A)**. Splenocytes from C57BL/6N mice were treated with 100ng/ml diABZI in culture for the indicated time. Dead cells were determined by PI and Annexin V staining. (**B).** Splenocytes from C57BL/6N mice were treated with indicated dose of diABZi in culture for 24hrs. Dead cells were determined by PI staining. **(C)**. Splenocytes from C57BL/6N mice were pre-treated with indicated small molecules, H151 (10µg/ml), C176 (1µM), MCC950 (1.25µM), 2-BP (20µM), GSK872 (312.5nM), Liproxstatin-1 (1µM), necrostatin-1 (10µM), VX-765 (1µM), Z-DEVD-FMK (25µM), Bx-795 (0.5µM) for 2hrs. diABZi (100ng/ml) was added in culture for another 24hrs. Dead cells were determined by PI staining. (**D)**. IFNβ was determined in the cell supernatant from **C** by ELISA. Data are representative of three independent experiments. Graphs represent the mean with error bars indication s.e.m. n=3∼5 mice/group. *p* values are determined by one-way ANOVA Tukey’s multiple comparison test (**A, D**) or unpaired student T-test (**C**). * p<0.05. n.s: not significant. n.d: not detected.

**Fig. S2.**
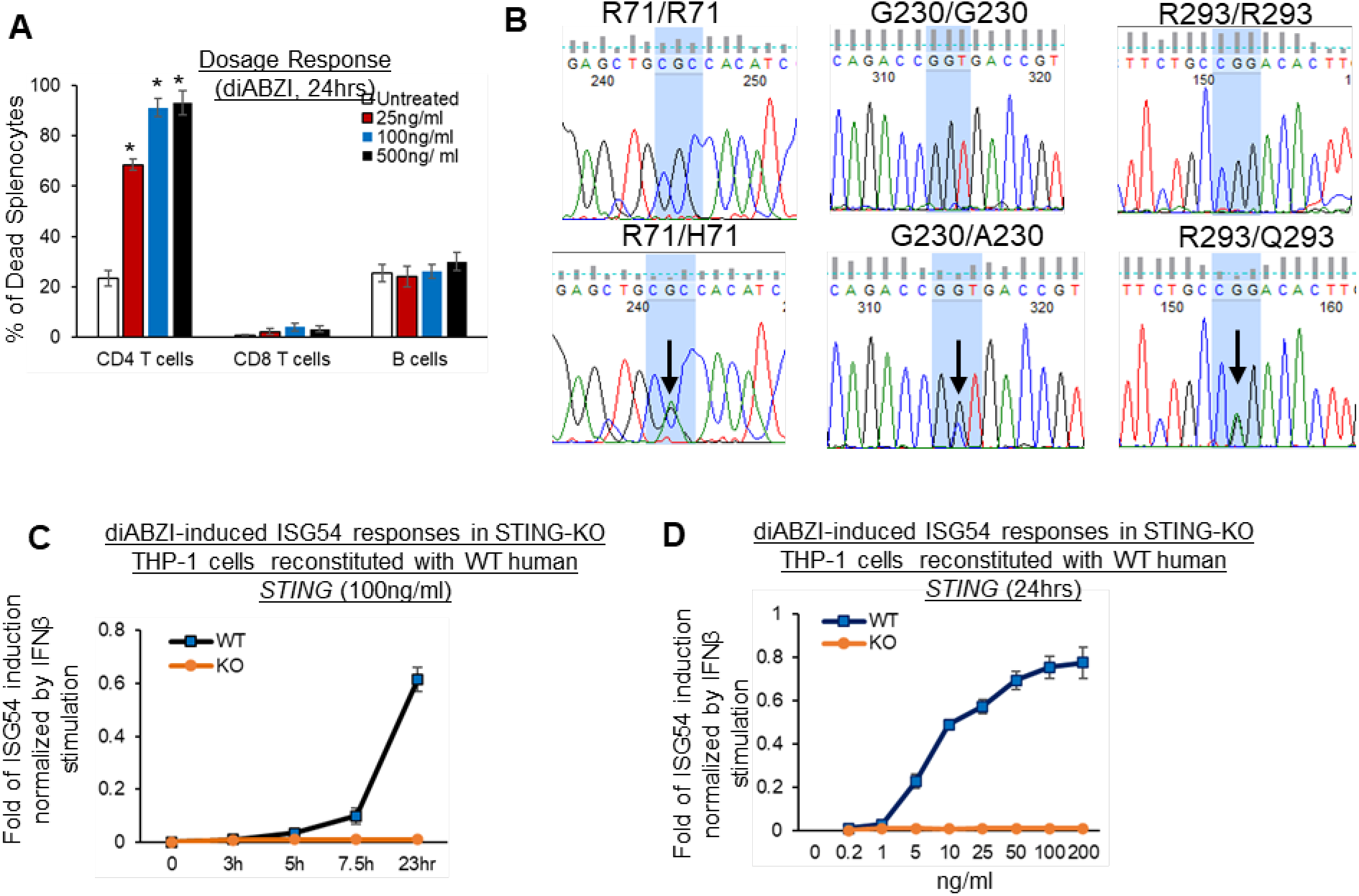
STING activation in primary human cells and THP-1 cells reconstituted with *WT* human *STING*. (**A**). Total human Lung cells from *WT/WT* individuals were treated with indicated dose of diABZi for 24hrs. Cell death in CD4, CD8 T cells and CD19 B cells were determined by PI staining. (**B).** Sequencing results of human *TMEM173* gene in *WT/WT* and *WT/HAQ* individuals. (**C-D)**. STING-KO THP-1 cells stably reconstituted with human *WT (R232)* were treated with indicated dose of diABZI for indicated time in culture. ISG-54 reporter luciferase activity was determined in cell supernatant and normalized to 10ng/ml IFNβ-stimulated ISG-54 luciferase activity. Data are representative of at least two independent experiments. Graphs represent the mean with error bars indication s.e.m. *p* values determined by one-way ANOVA Tukey’s multiple comparison test. * *p*<0.05, n.s: not significant.

**Fig. S3.**
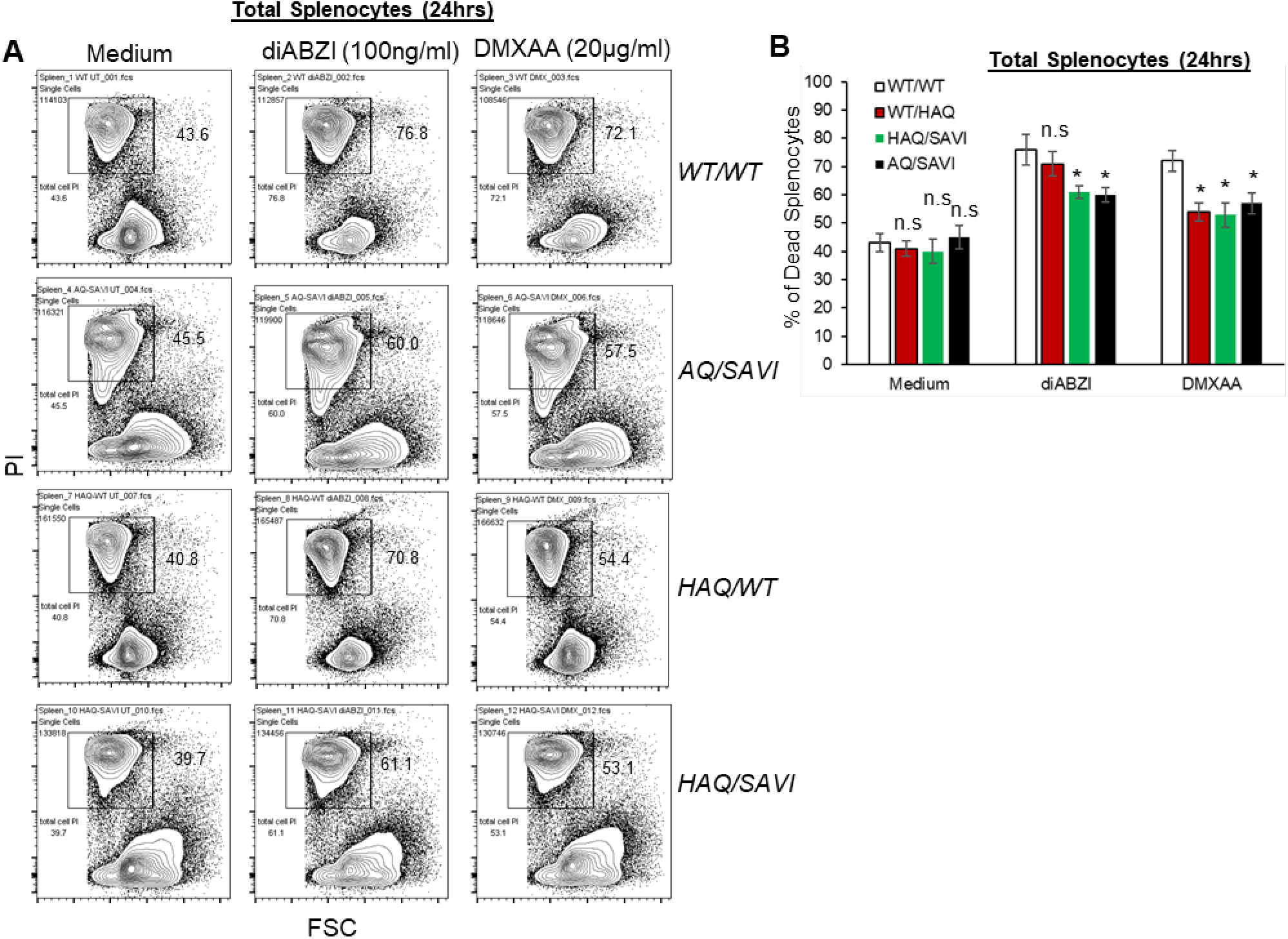
Splenocytes from HAQ/SAVI, AQ/SAVI partially resist to STING-activation-induced cell death ex vivo. **(A-B).** Flow cytometry of *HAQ/SAVI*, *AQ/SAVI*, *WT/WT* or *WT/HAQ* splenocytes treated with diABZI (100ng/ml) or DMXAA (20µg/ml) for 24hrs. Cell death was determined by PI staining. Data are representative of three independent experiments. Graphs represent the mean with error bars indication s.e.m. *p* values are determined by one-way ANOVA Tukey’s multiple comparison test. * p<0.05. n.s: not significant.

**Fig. S4.**
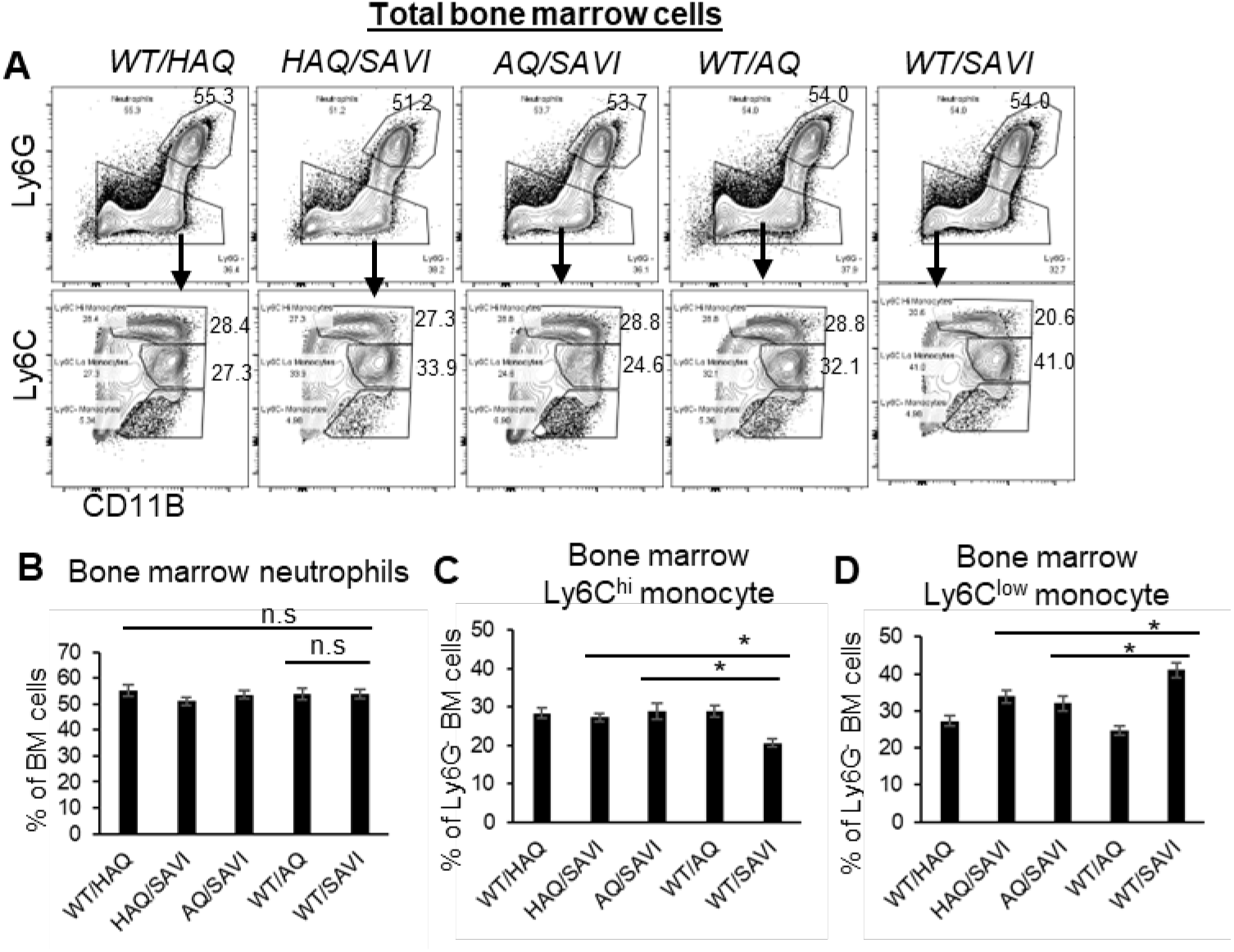
*HAQ* and *AQ* restore bone marrow monocytes in *SAVI (N153S)* mice. (**A-D**). Flow analysis of Ly6G^+^CD11B^+^ neutrophils and Ly6G^-^Ly6C^+^CD11B^+^ bone marrow monocytes from HAQ/SAVI, AQ/SAVI, WT/SAVI and their littermates. n=3∼5 mice/group. Data are representative of three independent experiments. Graphs represent the mean with error bars indication s.e.m. *P* values are determined by one-way ANOVA Tukey’s multiple comparison test. * *p*<0.05, n.s: not significant.

**Fig. S5.**
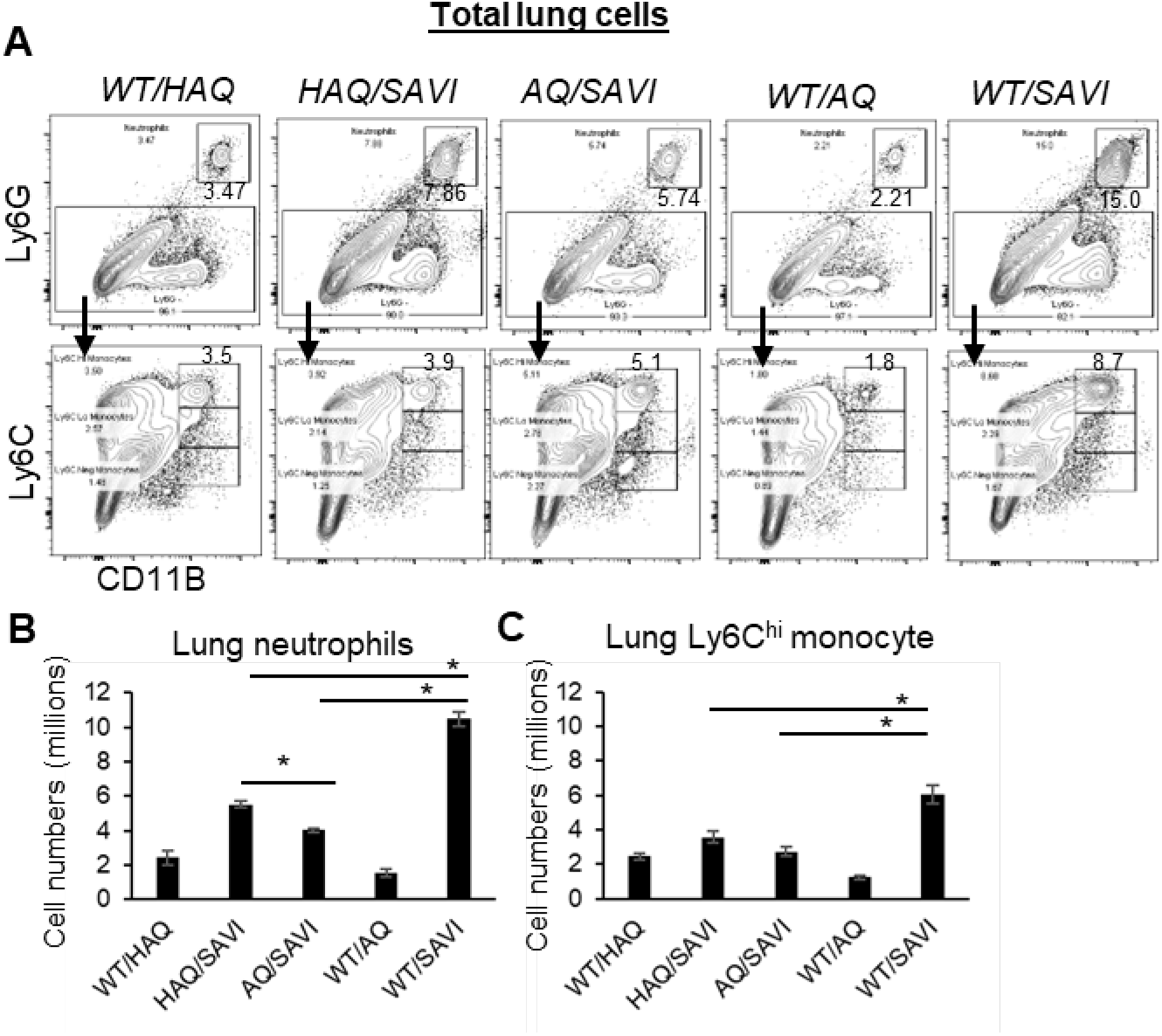
*HAQ, AQ* suppress lung myeloid cells infiltration in *SAVI(N153S)* mice. **(A-C)**. Flow analysis of lung Ly6G^+^CD11B^+^ neutrophils and Ly6G^-^Ly6C^+^CD11B^+^ monocytes from *HAQ/SAVI, AQ/SAVI, WT/SAVI and their littermates WT/HAQ, WT/AQ* mice. n=3∼5 mice/group. Data are representative of three independent experiments. Graphs represent the mean with error bars indication s.e.m. *P* values are determined by one-way ANOVA Tukey’s multiple comparison test. * *p*<0.05.

